# Augmented hip proprioception influences mediolateral foot placement during walking

**DOI:** 10.1101/2020.11.13.381665

**Authors:** Holly A. Knapp, Blaire A. Sobolewski, Jesse C. Dean

**Affiliations:** Medical University of South Carolina

**Keywords:** Biomechanics, Legged locomotion, Perturbation methods

## Abstract

Hip abductor proprioception contributes to the control of mediolateral foot placement, which varies with step-by-step fluctuations in pelvis dynamics. While prior work has used hip abductor vibration as a sensory perturbation to investigate this role of hip proprioception, we here tested whether time-varying vibration can predictably manipulate the relationship between pelvis dynamics and foot placement. We compared participants’ (n=32; divided into two groups of 16 with slightly different vibration control) gait behavior across four treadmill walking conditions: 1) No feedback; 2) Random feedback, with vibration unrelated to pelvis motion; 3) Augmented feedback, with vibration designed to evoke proprioceptive feedback paralleling the actual pelvis motion; 4) Disrupted feedback, with vibration designed to evoke proprioceptive feedback inversely related to pelvis motion. We hypothesized that the relationship between pelvis dynamics and foot placement would be strengthened by Augmented feedback but weakened by Disrupted feedback. For both participant groups, the strength of the relationship between pelvis dynamics at the start of a step and foot placement at the end of a step was significantly (p≤0.0002) influenced by the feedback condition. This metric was highest with Augmented feedback, but not significantly reduced with Disrupted feedback, partially supporting our hypotheses. Our approach to augmenting proprioceptive feedback during gait may have implications for clinical populations with a weakened relationship between pelvis motion and foot placement.

## I. Introduction

DURING human walking, pelvis dynamics predicts the mediolateral foot placement of the swing leg. Steps in which the pelvis has a large mediolateral displacement or velocity away from the stance foot end with more lateral foot placement relative to the pelvis, resulting in wider steps [1], [2]. The relationship between pelvis motion and foot placement is believed to be an important gait stabilization strategy [2], [3]; humans appear to take a wide enough step to avoid a lateral loss of balance, but not so wide as to cause excessive energy losses [4].

While step-by-step fluctuations in pelvis dynamics likely influence step width due to the body’s passive dynamics (e.g. the stance leg and trunk acting as an inverted pendulum), active control of the swing leg also plays a role. Larger mediolateral pelvis displacements and velocities are accompanied by stronger activation of the swing leg gluteus medius [5]. In turn, stronger gluteus medius activation is followed by more lateral foot placement of the swing leg, as would be predicted from this muscle’s action as a hip abductor [5]. The same pattern of adjusting gluteus medius activity and foot placement based on pelvis dynamics occurs when mediolateral perturbations are applied to the trunk or leg [5]–[7]. Additionally, manipulating the perception of body dynamics through visual [8] or vestibular [9] sensory perturbations elicits effects predicted by the dynamics-dependent active control of foot placement.

Our focus is on the potential of hip proprioceptive feedback to contribute to the relationship between pelvis motion and foot placement. The role of proprioceptive feedback in sensorimotor control can be probed using musculotendon vibration, a strong stimulus for activating primary muscle spindles and evoking the perception of musculotendon lengthening [10]. Vibration of the unilateral hip abductors during standing causes lateral sway away from the vibration [11], [12], as expected with the perception of a more adducted hip that would normally accompany lengthening of the vibrated muscle. In walking, perturbing proprioception through vibration of the stance hip abductors causes the swing foot to be placed more medially [12], [13]. This behavioral response is consistent with participants perceiving the pelvis to be closer mediolaterally to the stance foot (as would occur with a more adducted stance hip) and adjusting their active control to achieve a more medial foot placement. Conversely, vibration of the swing hip abductors causes the swing foot to be placed more laterally [12], likely a response to the evoked perception of the swing leg being overly adducted. The clearest effects of vibration are on mediolateral foot placement – the location of the swing foot relative to the pelvis – as the effects of vibration on stance leg motion are smaller and less consistent

In contrast with prior work that used hip abductor vibration as a discrete perturbation [11]–[13], we tested whether vibration could be controlled to manipulate the dynamics-dependent adjustments in foot placement. It is presently unclear whether this method can effectively augment available proprioceptive information during walking. If successful, such stimulation may allow individuals to achieve a stronger link between real-time pelvis dynamics and the foot placement control thought to be important for mediolateral balance.

The purpose of this study was to investigate whether targeted hip abductor vibration predictably influences the control of mediolateral foot placement during walking. Specifically, we applied vibration that could vary with each step, and took the form of four distinct feedback patterns: 1) No feedback, in which vibration was not applied; 2) Random feedback, in which vibration was unrelated to pelvis motion; 3) Augmented feedback, in which vibration scaled with the expected proprioceptive feedback from natural step-by-step variation in pelvis motion; 4) Disrupted feedback, in which vibration scaled inversely with the expected proprioceptive feedback. We hypothesized that Augmented feedback would strengthen the relationship between pelvis motion and foot placement, whereas Disrupted feedback would weaken this relationship.

## II. Methods

### A. Participants

16 young neurologically-intact participants (9 female / 7 male; age=23±2 yrs; height=176±8 cm; mass=77±20 kg; mean±s.d.) completed Experiment 1 of this study. A separate group of 16 young neurologically-intact participants (8 female / 8 male; age=23±1 yrs; height=174±11 cm; mass=74±14 kg) completed Experiment 2. Both Experiments 1 and 2 investigated the effects of hip abductor vibration on the control of mediolateral foot placement while walking. The experiments differed slightly in terms of how the vibration was controlled, providing insight into whether the effects of hip vibration were generalizable. All participants provided written informed consent using a form approved by the Medical University of South Carolina Institutional Review Board and consistent with the Declaration of Helsinki.

### B. Experimental Protocol

Participants performed a series of five treadmill walking trials at 1.2 m/s, wearing a harness attached to an overhead rail that did not support body weight, but would have prevented a fall in case of a loss of balance. An initial 10-minute accommodation trial allowed participants to become accustomed to walking on the treadmill. The remaining four 6-minute trials were performed in randomized order, and differed in terms of the delivered vibratory feedback: 1) No feedback; 2) Random feedback; 3) Augmented feedback; 4) Disrupted feedback. The control of Augmented feedback and Disrupted feedback differed between Experiment 1 and Experiment 2, as described in the Vibration Characteristics and Control section below.

### C. Data Collection and Processing

Active LED markers (PhaseSpace; San Leandro, CA, USA) were placed on the sacrum and bilateral heels. Marker position was sampled at 120 Hz, and low-pass filtered at 10 Hz. The start of each step was defined as when the ipsilateral heel velocity changed from posterior to anterior [14]. The end of each step was defined as when the contralateral heel velocity changed from posterior to anterior. At the start of each step, we calculated the mediolateral displacement of the sacrum with respect to the leading stance foot. We defined the positive mediolateral direction as toward the stepping leg.

### D. Vibration Characteristics and Control

Vibration was applied to the hip abductors using tactors (EMS^2^ tactors; Engineering Acoustics Inc.; Casselberry, FL, USA) secured over the bilateral gluteus medii, at the midpoint between the greater trochanter and the lateral iliac crest. These tactors consist of eccentrically weighted rotating motors in a plastic case, and are designed to produce vibration frequencies in a range that is excitatory for muscle spindles. In pilot testing, we found that both vibration frequency and displacement amplitude increased non-linearly with the applied control gain (Fig. 1). The frequency increased from 34-74 Hz, a range over which higher vibration frequencies evoke stronger primary muscle spindle responses [15]. The peak-to-peak displacement increased from 0.07-0.95 mm, with larger displacements again linked to stronger primary muscle spindle responses [16]. We focused our control on vibration frequency, as it continued to increase over a higher range of control gains.

**Fig. 1.**
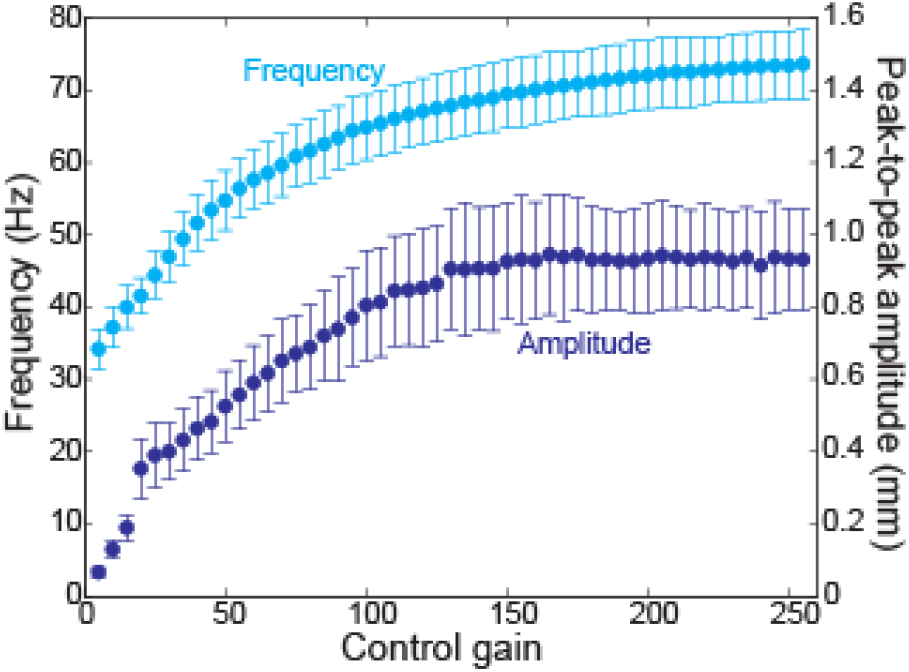
Vibration characteristics varied with control gain. Data points and error bars indicate means and 95% C.I. from pilot experiments (n=9) in which the tactors were securely strapped over the lateral aspect of seated participants’ thighs. Vibration characteristics were measured using a laser vibrometer (Optical Measurement Systems; Laguna Hills, CA, USA).

For Experiment 1, we first characterized each participant’s mean step period and the distribution of mediolateral pelvis displacement values at the start of the step for the final 50 steps (with each leg) of the accommodation trial. Across these 50 steps, we calculated the median pelvis displacement value, the 2^nd^ percentile value, and the 98^th^ percentile value (Fig. 2a). The vibration frequency in subsequent trials was chosen based on this Constant Distribution of displacement values, as detailed below. The remaining trials in Experiment 1 each corresponded to a different vibratory feedback condition. In the No feedback condition, no vibration was applied. For other trials, vibration was applied to either the stance or swing hip in a given step, turned on at the start of each step and turned off after a duration equal to the participant’s mean step period.

**Fig. 2.**
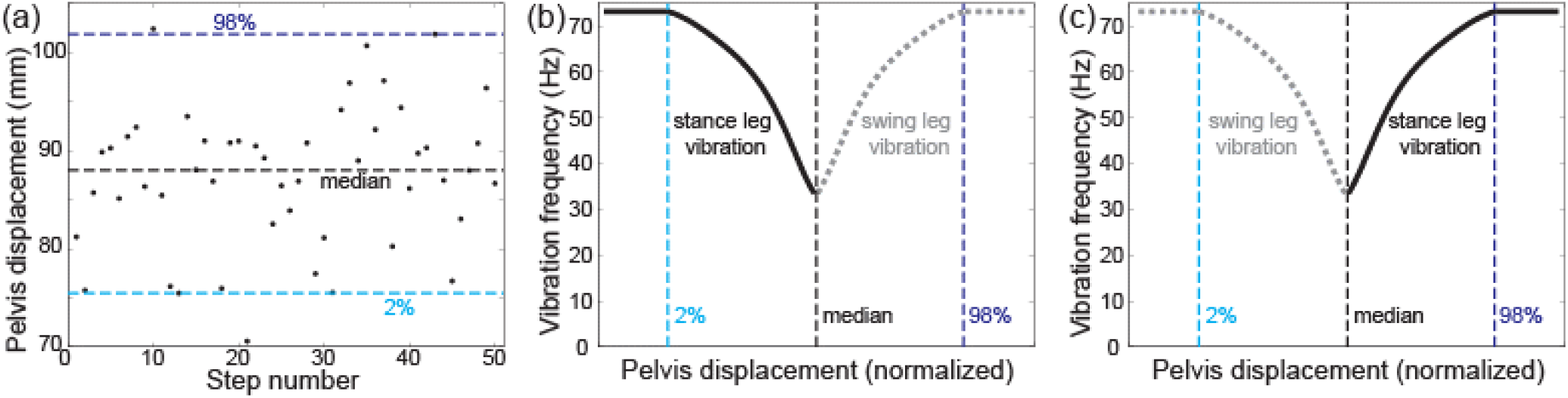
For some trials, vibration frequency was based on mediolateral pelvis displacement. We identified the median, 2%, and 98% pelvis displacement values for the final 50 steps of the accommodation trial, as illustrated for a single participant (a). With Augmented feedback (b), pelvis displacement was identified at the start of each step, and compared to the participant’s prior distribution of pelvis displacements. Displacements smaller than the median caused stance leg vibration, with higher vibration frequencies as the displacement approached the 2% value (and maximal for any smaller values). Displacements larger than the median caused swing leg vibration, with higher frequencies as the displacement approached the 98% value. With Disrupted feedback (c), the vibration side was reversed. The non-linear relationship in panels (b) and (c) produced a steeper frequency gradient for changes in displacement closer to the median. As we anticipated a near-normal distribution of values, ~68% of the frequency range is within an estimated standard deviation of the median, where we expected ~68% of the pelvis displacements.

In the Random feedback condition, the vibration was not linked to pelvis dynamics. Instead, the vibration side (stance or swing leg) was randomly chosen and the tactor control gain was randomly selected from a normal distribution.

In the Augmented feedback condition, vibration was applied to mimic sensory feedback that normally contributes to the control of mediolateral foot placement. For example, when the pelvis is close mediolaterally to the stance foot, humans tend to place their swing leg in a relatively medial location [1], [2]. Therefore, for pelvis displacements smaller than the median value, we applied stance leg vibration – which causes the foot to be placed more medially [12], [13]. In contrast, for pelvis displacements larger than the median value, we applied swing leg vibration – which causes more lateral foot placement [12], again matching the response that normally occurs with larger pelvis displacements [1], [2]. The applied vibration frequency scaled as illustrated in Fig. 2b; more extreme pelvis displacement values were accompanied by higher vibration frequencies.

In the Disrupted feedback condition, vibration was linked to pelvis displacement, but provided false information. The pattern of vibration was reversed from the Augmented feedback trial (Fig. 2c). Small pelvis displacements were accompanied by vibration that tends to evoke more lateral foot placement, while large pelvis displacements were accompanied by vibration that tends to evoke more medial foot placement.

From initial analysis of Experiment 1, we noted that despite our inclusion of a 10-minute accommodation trial, participants tended to walk with smaller mediolateral pelvis displacement values as the experiment progressed, possibly due to increased familiarity with treadmill walking. This shift biased our vibration control toward stance leg vibration in the Augmented feedback trial and swing leg vibration in the Disrupted feedback trial. Therefore, we performed an Experiment 2 to test whether our results were generalizable once this bias was addressed. Experiment 2 was structured identically to Experiment 1, except the vibration frequencies were based on an Updated Distribution of pelvis displacement values. This distribution was recalculated for each step, based on the most recent recorded 50 steps with this leg, accounting for potential gradual changes in the distribution of pelvis displacement values.

### E. Data Analysis and Statistics

Analyses were based on the first 250 steps from each of the four 6-minute trials, allowing comparisons across a consistent number of steps for each participant. Step length (SL) was calculated as the difference between the anterior position of the ipsilateral heel at the step end and the anterior position of the contralateral heel at the previous step end, accounting for treadmill speed. Step width (SW) was calculated as the mediolateral displacement between the ipsilateral heel marker at the step end and the contralateral heel marker at the step start. Mediolateral foot placement (FP) was calculated as the mediolateral displacement between the ipsilateral heel marker and the sacrum marker at the step end. Final mediolateral pelvis displacement (PD) was calculated as the mediolateral displacement between the sacrum marker at the step end and the contralateral heel marker at the step start. Therefore, step width is simply the sum of mediolateral foot placement (an estimate of swing leg positioning) and final mediolateral pelvis displacement (an estimate of stance leg positioning).

#### Distribution of vibration frequencies

For all trials with vibration, we identified the vibration side (stance or swing leg) and frequency for each step. We used a Wilcoxon rank sum test to determine whether the frequency distribution differed between the corresponding trials in Experiment 1 (Constant Distribution) and Experiment 2 (Updated Distribution). Stance leg vibration frequencies were assigned negative values and swing leg vibration frequencies were assigned positive values, allowing us to detect potential shifts in the overall distribution.

#### Within-trial comparison of stance and swing leg vibration

We here combined data from the Random feedback trials in Experiments 1 and 2, as the control of vibration in these trials did not differ between Experiments. For each participant, we used linear regressions to identify the coefficients for the following equations, for all steps with stance leg vibration:

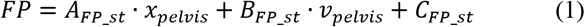

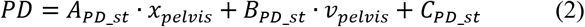

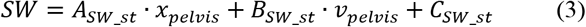

Here, x_pelvis_ is the mediolateral displacement between the pelvis and stance foot and v_pelvis_ is the mediolateral velocity of the pelvis at the start of the step. A, B, and C terms are the best fit coefficients, predicting mediolateral foot placement (FP), final pelvis displacement (PD), and step width (SW).

We performed identically-structured regressions to identify the coefficients for steps with swing leg vibration:

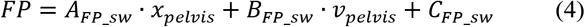

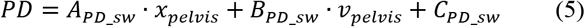

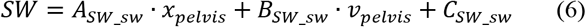

For each of the three corresponding regression equations, we performed paired t-tests to determine whether each of the regression coefficients differed between steps with stance leg vibration and steps with swing leg vibration.

#### Across-trial comparisons of vibration control methods

For the four feedback conditions (No feedback, Random feedback, Augmented feedback, Disrupted feedback), we quantified the relationship between pelvis dynamics throughout a step and the mediolateral body configuration at the end of the step. We resampled mediolateral pelvis displacement and velocity during each step to create 101-sample vectors, allowing comparisons across steps of variable periods [17]. For each normalized time point (0-100), we performed a linear regression in which the mediolateral pelvis displacement and velocity were used to predict mediolateral foot placement at the end of the step. The strength of this relationship was quantified as the R^2^ value from the regression. Our primary analysis focuses on the R^2^ value at the start of a step (step start R^2^_FP_), a measure providing insight into whether the within-step control of foot placement is influenced by vibration. Statistical analyses were performed separately for participants in Experiment 1 and Experiment 2, with a repeated-measures one-way ANOVA used to determine whether step start R^2^_FP_ differed significantly across the four feedback conditions. In the case of a significant main effect, Tukey-Kramer post-hoc tests were used to identify significant differences between individual conditions. Our design was not structured to directly test for differences between Experiment 1 and Experiment 2. Instead, we investigate whether consistent effects are observed with these control paradigms.

Secondarily, we investigated potential changes in the strength of the relationship between pelvis dynamics at the end of the step and mediolateral foot placement (step end R^2^_FP_), focusing on the moment when the new base of support was established. We also performed identically structured regressions to predict final mediolateral pelvis displacement based on pelvis dynamics at the start of the step (step start R^2^_PD_), and to predict step width based on pelvis dynamics at the start and end of a step (step start R^2^_SW_ and step end R^2^_SW_) for all four feedback conditions. For each feedback condition, we calculated each participant’s mean and standard deviation for step width, step length, and mediolateral foot placement. Potential effects of feedback condition on all of these secondary measures were investigated using the same repeated measures ANOVA statistical structure described above.

The main text focuses on R^2^ values as a measure of the strength of the relationship between pelvis dynamics and mediolateral stepping behavior. More detailed analyses of the specific contributions of mediolateral pelvis displacement and velocity are presented in the Supplementary Material.

#### Exploratory analysis of possible habituation to vibration

To investigate whether the effects of vibration changed with repeated exposure, we divided the 250 steps from each feedback condition into five blocks of 50 steps. Our analysis focused on our primary outcome measure (step start R^2^_FP_), and we combined data from Experiments 1 and 2 given the similarities between the observed results. We performed a repeated measures two-way ANOVA, with block number (1-5) and feedback condition as independent variables.

For all statistical analyses, p-values less than 0.05 were interpreted as significant.

## III. Results

### Distribution of vibration frequencies

The distribution of vibration frequencies differed across the Random, Augmented, and Disrupted feedback conditions. For the Random condition, vibration was equally applied to the stance and swing legs, albeit not equally distributed across the frequency range due to non-linearities in the relationship between control gain and frequency. This distribution was similar for Experiment 1 (Fig. 3a) and Experiment 2 (Fig. 3d), with no significant difference between the median vibration frequencies (p=0.51). In contrast, the frequency distribution differed between the Augmented conditions in Experiment 1 (Fig. 3b) and Experiment 2 (Fig. 3e). When the relationship between pelvis displacement and vibration frequency was held constant (Experiment 1), the distribution was biased toward vibration of the stance leg (median=54 Hz stance leg vibration). Updating this relationship while participants walked (Experiment 2) significantly (p<0.0001) reduced the magnitude of this bias (median=36 Hz stance leg vibration). Similarly, the frequency distribution differed between the Disrupted conditions in Experiment 1 (Fig. 3c; median=53 Hz swing leg vibration) and Experiment 2 (Fig. 3f; median=39 Hz swing leg vibration), with the bias toward swing leg vibration significantly (p<0.0001) reduced when the control relationship was updated.

**Fig. 3.**
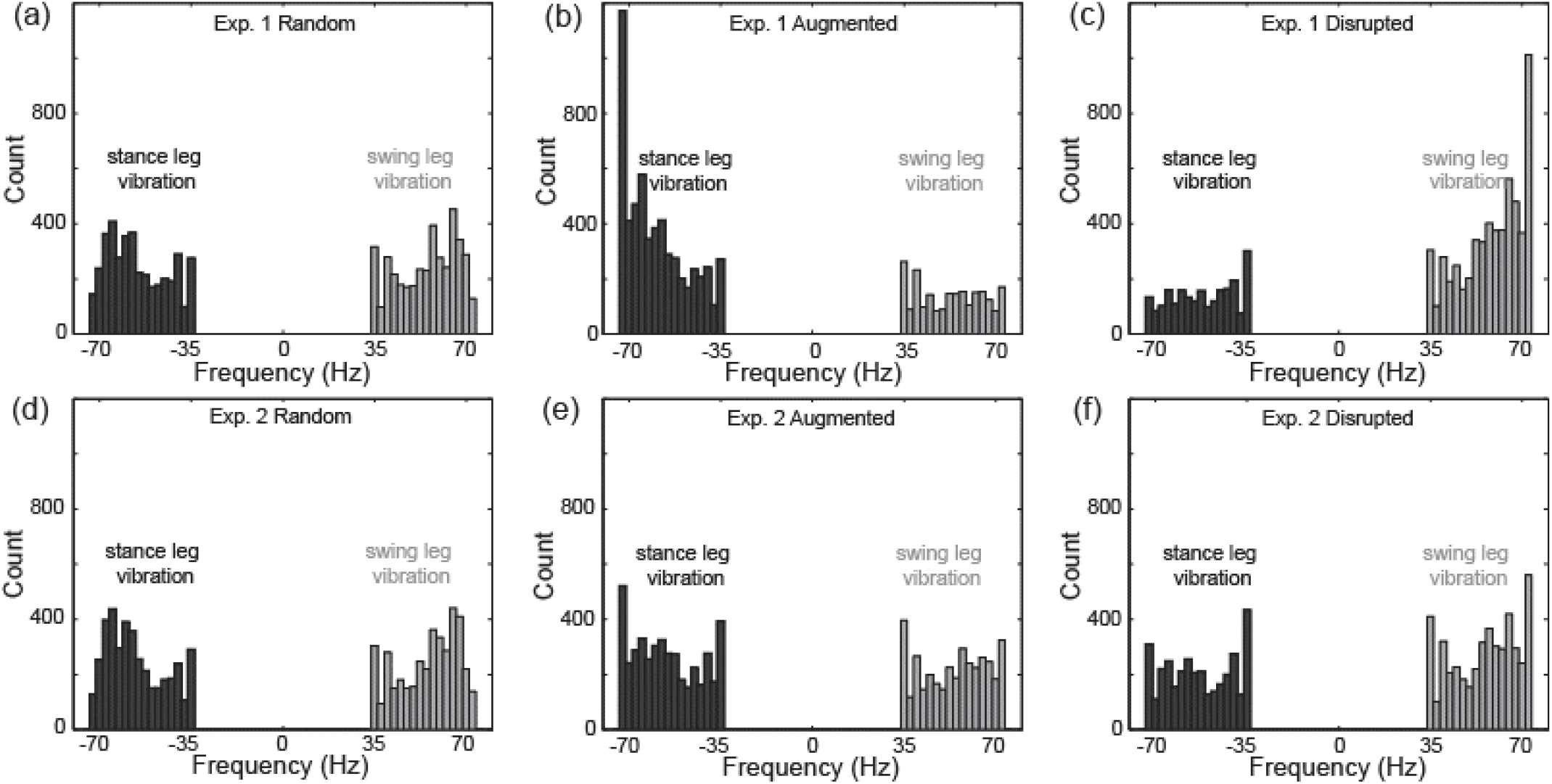
The distribution of vibration frequencies differed across conditions and Experiments, as illustrated for Experiment 1 in the top row of panels (a-c) and for Experiment 2 in the bottom row (d-f). For ease of illustration, stance leg vibration frequencies are plotted as negative values.

### Within-trial comparison of stance and swing leg vibration

During the Random condition, stance leg vibration and swing leg vibration differentially affected mediolateral foot placement and step width. Table I presents the coefficient values from regression equations 1-6 (detailed in the Methods). In no case did the regression coefficient for mediolateral displacement or velocity differ between stance and swing leg vibration (p≥0.25 for all comparisons), indicating that step-by-step fluctuations in pelvis dynamics consistently predict fluctuations in body configuration at the end of the step. However, steps with swing leg vibration had significantly larger offset values than those with stance leg vibration for both mediolateral foot placement (p=0.012) and step width (p=0.004), indicating that swing leg vibration caused more lateral foot placement and wider steps. No such significant difference was observed for final pelvis displacement (p=0.13).

**TABLE I.** Regression coefficients

|  |  | Stance leg vibration | Swing leg vibration | Stance vs. swing p-value |
| --- | --- | --- | --- | --- |
| Foot placement predictors | A<sub>FP</sub> | 0.48±0.04 | 0.47±0.03 | 0.44 |
|  | B<sub>FP</sub> | 0.070±0.012 | 0.071±0.011 | 0.85 |
|  | C<sub>FP</sub> | 38.5±6.6 | 40.5±6.6 | **0.012** |
| Pelvis displacement predictors | A<sub>PD</sub> | 0.81±0.03 | 0.80±0.02 | 0.56 |
|  | B<sub>PD</sub> | 0.069±0.011 | 0.073±0.010 | 0.25 |
|  | C<sub>PD</sub> | 28.9±3.4 | 29.7±3.5 | 0.13 |
| Step width predictors | A<sub>SW</sub> | 1.29±0.05 | 1.27±0.04 | 0.38 |
|  | B<sub>SW</sub> | 0.14±0.02 | 0.14±0.02 | 0.53 |
|  | C<sub>SW</sub> | 67.4±7.9 | 70.2±7.8 | **0.004** |

Coefficient values for linear regressions used to predict mediolateral foot placement, final pelvis displacement, and step width. Values are presented as means ± 95% C.I.

### Across-trial comparisons of vibration control methods

The effects of hip vibration varied across feedback conditions. For illustrative purposes, Fig. 4 depicts the R^2^ values calculated throughout a step – showing the proportion of the variability in mediolateral foot placement (Fig. 4a, 4d), final pelvis displacement (Fig. 4b, 4e), and step width (Fig. 4c, 4f) that can be predicted from pelvis dynamics.

**Fig. 4.**
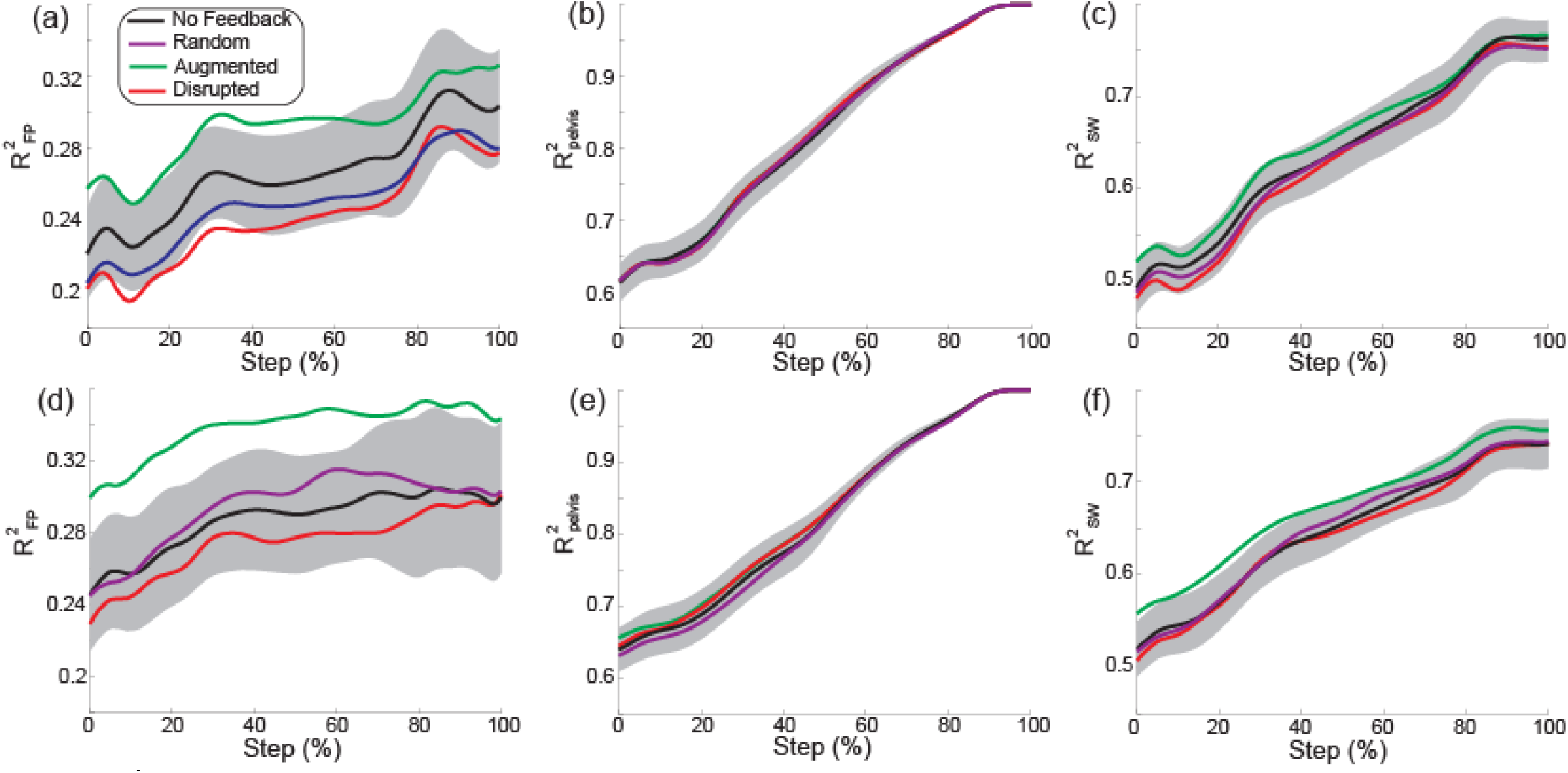
The R^2^ values calculated based on pelvis dynamics throughout a step (step start = 0%; step end = 100%) are illustrated for each of our metrics. The top row (a-c) presents results from Experiment 1, while the bottom row presents results from Experiment 2. From left to right, we illustrate the proportion of mediolateral foot placement (a, d), final mediolateral pelvis displacement (b, e), and step width (c, f) that is predicted from pelvis dynamics. Solid lines represent mean values, and the shaded gray area represents the 95% C.I. for the No Feedback condition. Error bars are not provided for the other conditions due to excessive overlap, but variability in the effects of vibration is illustrated in Fig. 5 below.

Our primary outcome measure (step start R^2^_FP_) varied significantly across conditions (Fig. 5a) for both Experiment 1 (p=0.0002) and Experiment 2 (p<0.0001). This metric was highest with Augmented feedback, but did not differ significantly across the remaining conditions. Step end R^2^_FP_ also varied significantly across conditions (Fig. 5b) for Experiment 1 (p=0.002) and Experiment 2 (p=0.017), although with less consistent significant differences between the Augmented feedback condition and other conditions.

**Fig. 5.**
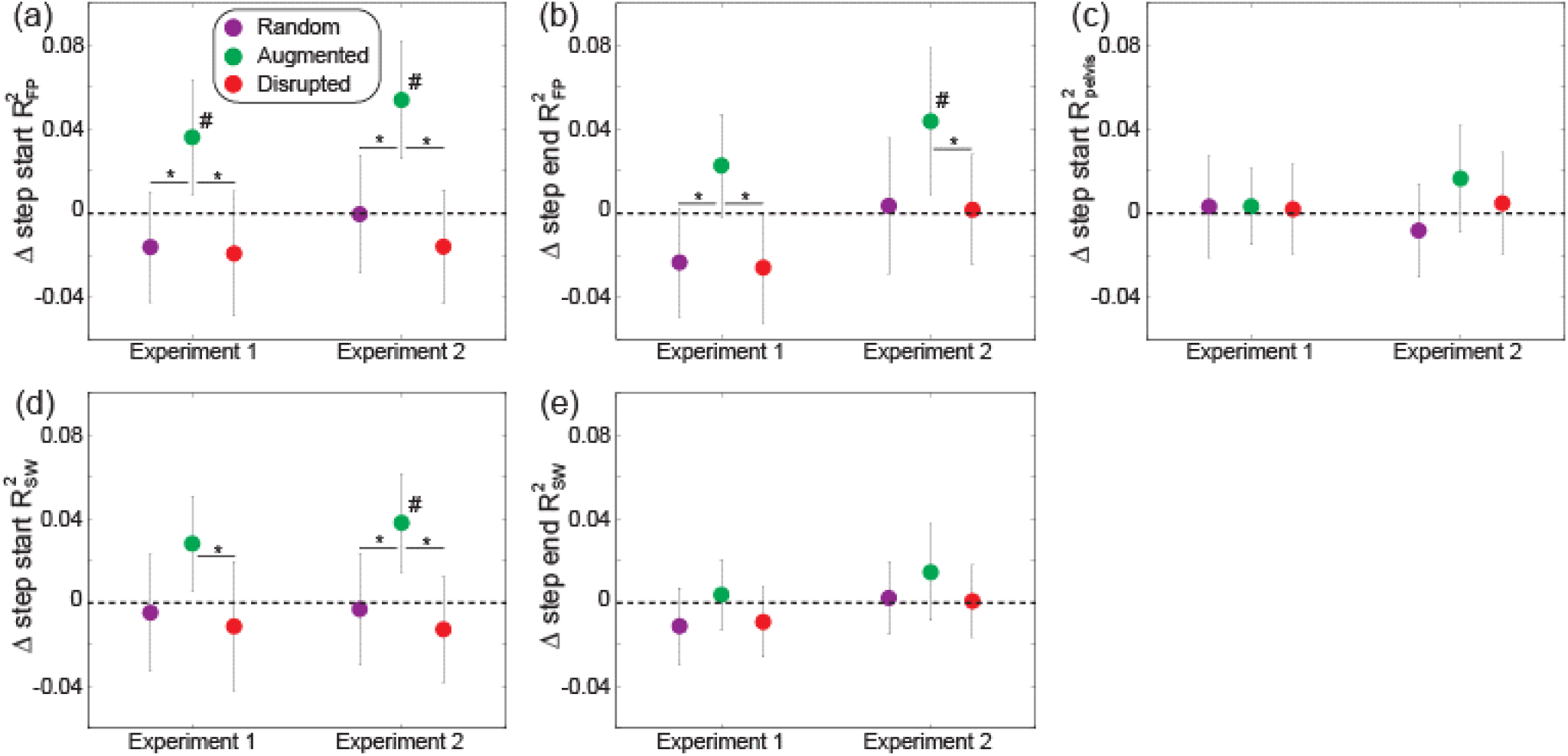
Several metrics of mediolateral gait behavior varied across feedback conditions. In all panels, we illustrate the change in the R^2^ metrics relative to the No feedback condition, to account for individual variability and better focus on the effects of vibration. Specifically, we illustrate the change in step start R^2^_FP_(a), step end R^2^_FP_(b), step start R^2^_PD_(c), step start R^2^_SW_(d), and step end R^2^_SW_(e). Data points indicate means and error bars indicate 95% C.I. Asterisks (*) indicate significant post-hoc differences between the indicated conditions, and pound signs (#) indicate significant post-hoc differences from the No Feedback condition.

Unlike mediolateral foot placement, the relationship between pelvis dynamics at the start of a step and the final pelvis displacement was not affected by vibration. Specifically, step start R^2^_PD_ did not vary significantly across conditions (Fig. 5c) for either Experiment 1 (p=0.99) or Experiment 2 (p=0.19).

The relationship between pelvis dynamics and step width was more equivocally influenced by hip vibration than foot placement. Step start R^2^_SW_ varied significantly across feedback conditions for both Experiment 1 (p=0.030) and Experiment 2 (p=0.0005), with the largest values with Augmented feedback (Fig. 5d). However, step end R^2^_SW_ did not vary significantly across conditions (Fig. 5e) for either Experiment 1 (p=0.23) or Experiment 2 (p=0.41).

None of the more traditional measures of gait mechanics varied significantly across conditions, as presented in Table II.

**TABLE II.** Gait spatial metrics

| Metric (mm) | Experiment 1 |  |  |  |  | Experiment 2 |  |  |  |  |
| --- | --- | --- | --- | --- | --- | --- | --- | --- | --- | --- |
|  | None | Random | Augment. | Disrupt. | P-value | None | Random | Augment. | Disrupt. | P-value |
| Mean SW | 120±11 | 118±11 | 121±11 | 122±12 | 0.10 | 125±10 | 127±10 | 127±11 | 123±10 | 0.18 |
| Mean SL | 677±11 | 676±11 | 677±11 | 676±12 | 0.82 | 663±14 | 662±15 | 664±15 | 663±14 | 0.74 |
| Mean FP | 57±9 | 56±8 | 58±9 | 58±9 | 0.29 | 57±7 | 59±7 | 58±8 | 56±7 | 0.20 |
| Std. SW | 18±1 | 17±1 | 18±1 | 17±1 | 0.15 | 19±1 | 18±1 | 19±2 | 18±2 | 0.32 |
| Std. SL | 13±1 | 13±1 | 13±1 | 14±2 | 0.12 | 14±1 | 14±1 | 13±1 | 13±1 | 0.23 |
| Std. FP | 10±1 | 10±1 | 10±1 | 10±1 | 0.19 | 11±1 | 11±1 | 11±1 | 11±1 | 0.22 |

Mean and variability of gait spatial metrics (step width [SW], step length [SL], and foot placement [FP]) across the four feedback conditions and the two experiments. Values are presented as means ± 95% C.I.

### Exploratory analysis of possible habituation to vibration

No direct evidence for habituation was observed (Fig. 6). Consistent with the primary results presented above, feedback condition had a significant main effect on step start R^2^_FP_ (p<0.0001), with the highest values for the Augmented condition. However, no significant main effect of block number (p=0.19) or an interaction between condition and block number (p=0.87) was detected.

**Fig. 6.**
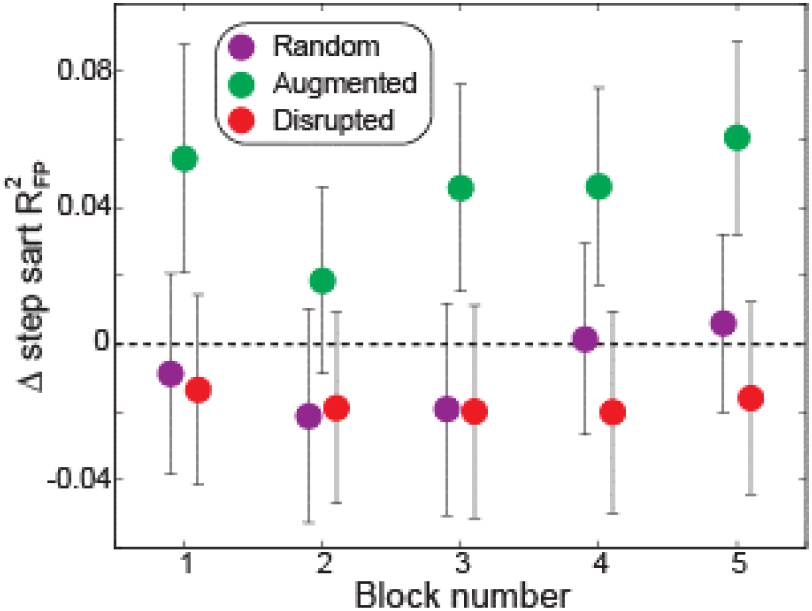
Our primary metric (step start R^2^_FP_) did not change significantly across 50-step blocks. Data points indicate means and error bars indicate 95% C.I.

## IV. Discussion

Humans actively control mediolateral foot placement in response to fluctuations in body dynamics, with hip proprioceptive feedback contributing to this control. We found that augmenting proprioceptive feedback using scaled vibration strengthened the relationship between pelvis dynamics and foot placement, as hypothesized. In contrast, using similar methods to disrupt proprioceptive feedback did not significantly weaken this relationship, contradicting our hypothesis. While we tested two control methods for delivering these patterns of vibration (based on a Constant Distribution or Updated Distribution of pelvis displacements), our discussion will focus on the consistent results observed with both of these control methods.

Overall, the relationship between pelvis dynamics and stepping behavior was consistent with previous work. Larger mediolateral pelvis displacements or velocities away from the stance foot predicted that the step would end with more lateral foot placement, larger mediolateral pelvis displacements, and wider steps [1], [2], [17]. These relationships were apparent from pelvis dynamics at the start of a step, and generally became stronger over the course of the step.

The effects of Random feedback were qualitatively similar to those observed in previous experiments in which the applied vibration was not linked to the actual pelvis motion [12], [13]. The effects of step-by-step fluctuations in pelvis dynamics were essentially identical for steps with either stance leg or swing leg vibration, indicating that the important role of pelvis motion was retained. However, in comparison to steps with stance leg vibration, swing leg vibration tended to produce more lateral foot placement and wider steps (by ~2-3 mm). Despite this differential effect of stance and swing leg vibration, the overall strength of the relationship between pelvis dynamics and stepping behavior (quantified with R^2^ values) was not significantly lower for the Random feedback condition than the No feedback condition. Speculatively, perhaps the sensory information provided by the Random hip vibration was recognized by the nervous system as unreliable, as the elicited proprioceptive signal likely contradicts information from other sensory sources (visual, vestibular, foot cutaneous) that contribute to the overall perception of the body’s mechanical state [8], [9], [18]. Such unreliable sensory information can be downweighted by the nervous system [19], possibly reducing the effects of Random feedback.

Augmented hip proprioceptive feedback affected the dynamics-dependent control of foot placement, but not the mediolateral motion of the pelvis - which may be strongly influenced by inverted pendulum mechanics. The relationship between pelvis dynamics (predominantly pelvis displacement; see Supplementary Material) throughout a step and mediolateral foot placement was strengthened for the Augmented feedback condition. Essentially, applying scaled stance leg vibration when the pelvis was close to the stance foot increased the likelihood that the swing leg trajectory would be adjusted to achieve a relatively medial foot placement, while applying scaled swing leg vibration when the pelvis was far from the stance foot increased the likelihood of a relatively lateral foot placement. This result is consistent with the previously observed effects of hip abductor vibration during walking [12], [13], and coincides with the typical foot placement strategy thought to be important for mediolateral balance [2]. Unlike with mediolateral foot placement (an estimate of swing leg positioning), Augmented feedback did not affect the relationship between pelvis dynamics throughout a step and pelvis displacement at the end of a step (an estimate of stance leg and trunk positioning). The clearer effect of hip vibration on swing leg motion is consistent with prior work [12], and could perhaps be due to the lower mechanical demands of repositioning the swing leg compared to controlling the motion of the more massive stance leg and trunk [20].

While the effects of Augmented feedback demonstrate the potential of manipulating step-by-step foot placement through targeted sensory stimulation, the present results do not allow us to differentiate between potential mechanisms underlying these effects. Sensory augmentation is often considered to complement information available from natural sensory signals, which may be compromised by inherent sensory noise [21]. Such augmented information may allow individuals to form a more accurate perception of their body’s dynamic state [22]. In the present work, the scaled stance leg vibration may increase the intensity of the proprioceptive signal from the stance hip abductors, thus creating a stronger perception of a small displacement between the pelvis and stance foot, and leading to swing leg actuation to produce more medial foot placement. In contrast, scaled swing leg vibration may create a stronger perception of an overly adducted swing leg, causing swing leg actuation to produce more lateral foot placement. This explanation is speculative, as we did not directly assess perception. Alternatively, perhaps the changes in foot placement were driven by simpler reflexive or spinal circuits known to play a role in locomotion [23]. Future work will be required to provide insight into the relative role of several mechanisms of sensory augmentation that could feasibly underlie improvements in human balance performance, as recently reviewed [24].

Unlike with Augmented feedback, Disrupted feedback did not significantly alter the relationship between pelvis dynamics and foot placement (or other gait metrics), contradicting our hypothesis. The relative ineffectiveness of Disrupted feedback at altering stepping behavior may be due to the mechanism of downweighting unreliable feedback cited above [19]. These limited effects indicate that participants were not simply using the applied vibration as a “cue” to adjust their gait pattern, a form of sensory addition explored in previous work [25]. In theory, the Disrupted feedback provided the nervous system with just as much information about the dynamic state of the pelvis as the Augmented feedback – simply switching the leg side to which this feedback was delivered. Participants did not use this information to strengthen the relationship between pelvis dynamics and foot placement, suggesting a benefit of applying feedback through natural sensory pathways (e.g. amplifying an afferent signal of stance leg hip abductor stretch when this is truly what is occurring mechanically) rather than requiring participants to discover a new linkage between novel afferent feedback and body dynamics [26].

The present results do not provide evidence for either habituation or adaptation with sustained exposure to hip vibration. Specifically, we did not observe a reduction in the effects of vibration over five consecutive 50-step blocks, as would occur if participants gradually habituated to ignore the afferent signal produced by vibration [27], [28]. Similarly, the effects of vibration did not increase over time, as may occur if participants only gradually adapt their sensorimotor control to take advantage of the information provided by vibration. While the lack of a change in the effects of vibration over time is consistent with prior work [13], our results don’t provide insight into habituation or adaptation that may occur over shorter (<50 steps) or longer (>250 steps) time frames.

While our results suggest that Augmented hip proprioceptive feedback can strengthen the relationship between pelvis dynamics and foot placement, several important limitations should be noted. The patterns of vibration used in this study reflect only a small subset of possible options. For example, vibration frequency could instead scale linearly with pelvis displacement (resulting in a more normal distribution of frequency values on each side of Fig. 3), vibration frequency could be updated continuously during a step as the pelvis moves rather than being changed once per step, or the stimulation intensity could be faded over time to test whether any changes in gait behavior are retained [29]. This study was also not designed to directly compare the effects of basing the vibration control on a Constant Distribution (Experiment 1) or Updated Distribution (Experiment 2) of pelvis displacement values, although the results appear quite similar. Our focus on hip proprioceptive feedback ignored other sensory sources that could also influence foot placement behavior (e.g. visual, vestibular, cutaneous) [8], [9], [18], and could conceivably be augmented using similar methods. Finally, this study did not directly investigate whether participants updated a learned relationship between perceived body dynamics and target foot placement location. Such an adjustment in this sensorimotor relationship may be required in order for the augmented proprioceptive feedback to strengthen the relationship between pelvis dynamics and foot placement without changing the overall distribution (mean and standard deviation) of foot placement locations, as observed here.

In the longer term, the application of Augmented hip proprioceptive feedback during walking may have clinical implications. Individuals with chronic stroke and poor walking balance exhibit a weakened relationship between pelvis dynamics, paretic swing leg hip abductor activation, and paretic mediolateral foot placement location [30]. Augmenting the available sensory feedback may allow more accurate adjustments of paretic foot placement based on body dynamics, with potential benefits for walking balance. As our sensory augmentation approach does not require participants to consciously attend to the vibration, it may have lower cognitive demands than more traditional “cueing” strategies [26], and may be particularly useful for patient populations in whom walking is already cognitively demanding.

In conclusion, a novel method of augmenting hip proprioceptive feedback successfully strengthened the relationship between pelvis dynamics and mediolateral foot placement, although the underlying mechanism is unclear. Disrupting proprioceptive feedback did not produce significant negative effects, possibly due to the availability of other sensory sources providing information about the body’s dynamic state. This approach may be applicable in clinical settings for individuals with walking balance deficits.

