## Supplementary Material for "Augmented hip proprioception influences mediolateral foot placement during walking"

THIS study used $R^2$ values calculated from linear regressions to quantify the strength of the relationship between pelvis dynamics and mediolateral stepping behavior. We calculated the proportion of the variation in mediolateral foot placement, final pelvis displacement, and step width that was predicted by the combination of mediolateral pelvis displacement and velocity. Here, we instead use partial correlations to focus on the separate contributions of pelvis displacement and pelvis velocity to the observed stepping behavior. The 10 quantified partial correlation metrics are listed in Table SI with corresponding brief descriptions.

As in the main text, statistical comparisons were performed separately for Experiment 1 and Experiment 2. We used repeated measures one-way ANOVA to determine whether each partial correlation metric differed across the four feedback conditions. In the case of a significant effect, we used Tukey-Kramer post-hoc tests to identify significant differences between individual conditions. For all comparisons, p-values less than 0.05 were interpreted as significant.

The partial correlation between mediolateral pelvis displacement and several metrics of stepping behavior differed across feedback conditions, paralleling our main text results investigating $R^2$ magnitude. Step start $\rho_{disp\_FP}$ varied significantly across conditions (Fig. S1a) for both Experiment 1 ($p=0.0002$) and Experiment 2 ($p<0.0001$), and was highest for the Augmented feedback condition. Step end $\rho_{disp\_FP}$ also varied significantly across conditions (Fig. S1b) for both Experiment 1 ($p=0.02$) and Experiment 2 ($p=0.0005$), and was highest with Augmented feedback. Step start $\rho_{disp\_PD}$ did not vary across conditions (Fig. S1c) for either Experiment 1 ($p=0.97$) or Experiment 2 ($p=0.17$). Step start $\rho_{disp\_SW}$ varied significantly across conditions (Fig. S1d) for both Experiment 1 ($p=0.049$) and Experiment 2 ($p=0.001$), and was highest for the Augmented feedback condition. Step end $\rho_{disp\_SW}$ did not vary across conditions (Fig. S1e) for either Experiment 1 ($p=0.29$) or Experiment 2 ($p=0.25$).

In contrast to our observations with pelvis displacement, none of the partial correlations focused on pelvis velocity varied

TABLE SI
PARTIAL CORRELATION METRICS

| Metric | Description |
|---|---|
| step start $\rho_{disp\_FP}$ | Partial correlation between mediolateral pelvis displacement at the start of a step and mediolateral foot placement at the end of the step, accounting for mediolateral pelvis velocity at the start of the step |
| step end $\rho_{disp\_FP}$ | Partial correlation between mediolateral pelvis displacement at the end of a step and mediolateral foot placement at the end of the step, accounting for mediolateral pelvis velocity at the end of the step |
| step start $\rho_{disp\_PD}$ | Partial correlation between mediolateral pelvis displacement at the start of a step and mediolateral pelvis displacement at the end of the step, accounting for mediolateral pelvis velocity at the start of the step |
| step start $\rho_{disp\_SW}$ | Partial correlation between mediolateral pelvis displacement at the start of a step and step width at the end of the step, accounting for mediolateral pelvis velocity at the start of the step |
| step end $\rho_{disp\_SW}$ | Partial correlation between mediolateral pelvis displacement at the end of a step and step width at the end of the step, accounting for mediolateral pelvis velocity at the end of the step |
| step start $\rho_{vel\_FP}$ | Partial correlation between mediolateral pelvis velocity at the start of a step and mediolateral foot placement at the end of the step, accounting for mediolateral pelvis displacement at the start of the step |
| step end $\rho_{vel\_FP}$ | Partial correlation between mediolateral pelvis velocity at the end of a step and mediolateral foot placement at the end of the step, accounting for mediolateral pelvis displacement at the end of the step |
| step start $\rho_{vel\_PD}$ | Partial correlation between mediolateral pelvis velocity at the start of a step and mediolateral pelvis displacement at the end of the step, accounting for mediolateral pelvis displacement at the start of the step |
| step start $\rho_{vel\_SW}$ | Partial correlation between mediolateral pelvis velocity at the start of a step and step width at the end of the step, accounting for mediolateral pelvis displacement at the start of the step |
| step end $\rho_{vel\_SW}$ | Partial correlation between mediolateral pelvis velocity at the end of a step and step width at the end of the step, accounting for mediolateral pelvis displacement at the end of the step |

Definitions of various partial correlation values used to quantify the contributions of pelvis displacement and velocity to stepping behavior significantly across feedback conditions ($p \geq 0.14$ for all comparisons). Values are illustrated in Figure S2 for visual comparison.

The present results indicate that the changes in $R^2$ magnitude described in the main text are predominantly due to a change in the step-by-step relationship between pelvis displacement and stepping behavior, and not to a change in the relationship between pelvis velocity and stepping behavior. The relative importance of pelvis displacement is likely caused in part by our choice to control vibration intensity based on pelvis displacement, not pelvis velocity.

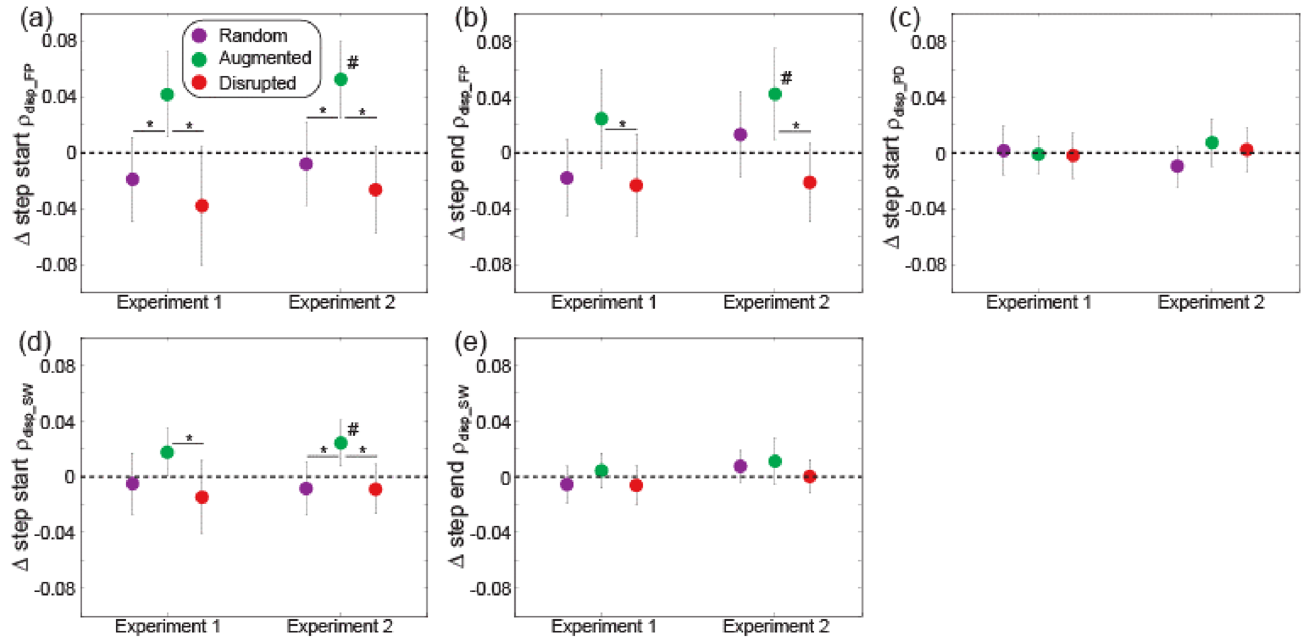


Fig. S1. Several metrics based on the partial correlation with mediolateral pelvis displacement varied across feedback conditions. As in the main text, we illustrate the change in the partial correlation metrics relative to the No feedback condition. We illustrate the change in step start $\rho_{disp\_FP}$ (a), step end $\rho_{disp\_FP}$ (b), step start $\rho_{disp\_PD}$ (c), step start $\rho_{disp\_SW}$ (d), and step end $\rho_{disp\_SW}$ (e). Data points indicate means and error bars indicate 95% C.I. Asterisks (*) indicate significant post-hoc differences between the indicated conditions, and pound signs (#) indicate significant post-hoc differences from the No feedback condition.

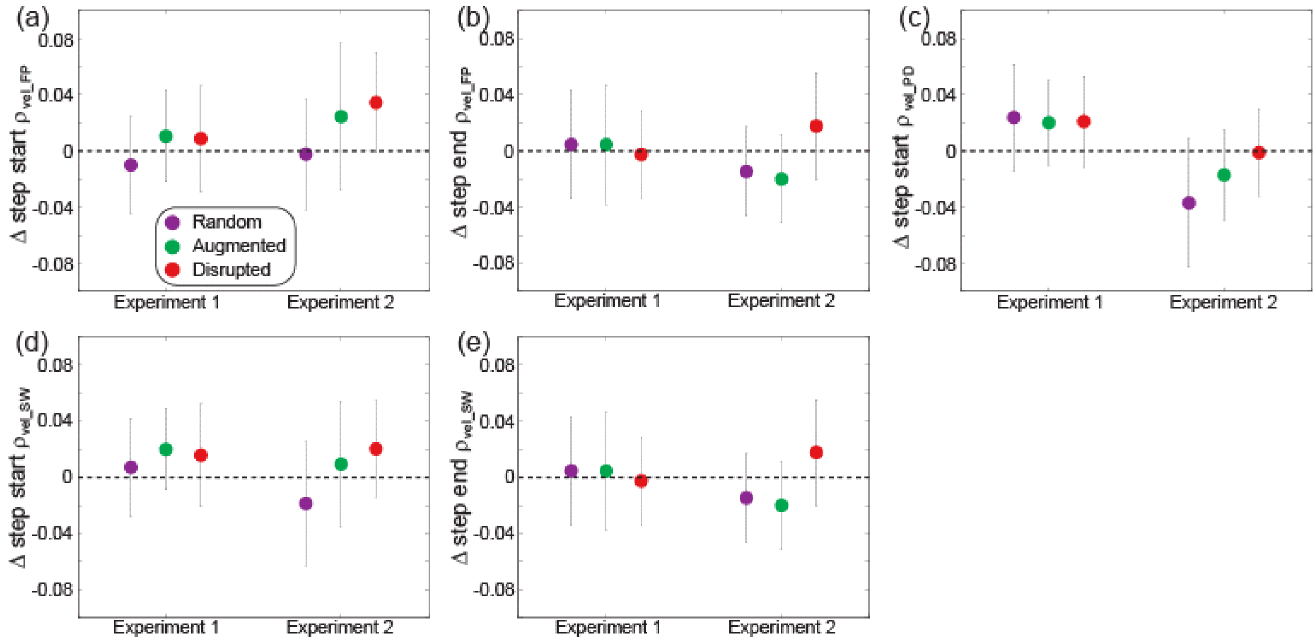


Fig. S2. Metrics based on the partial correlation with mediolateral pelvis velocity did not vary across feedback conditions, as illustrated relative to the No feedback condition. We illustrate the change in step start $\rho_{vel\_FP}$ (a), step end $\rho_{vel\_FP}$ (b), step start $\rho_{vel\_PD}$ (c), step start $\rho_{vel\_SW}$ (d), and step end $\rho_{vel\_SW}$ (e). Data points indicate means and error bars indicate 95% C.I.
